# Sequential 3D OrbiSIMS and LESA-MS/MS-based metabolomics for prediction of brain tumor relapse from sample-limited primary tissue archives

**DOI:** 10.1101/2020.07.15.182071

**Authors:** Joris Meurs, David J. Scurr, Arockia Lourdusamy, Lisa C.D. Storer, Richard G. Grundy, Morgan R. Alexander, Ruman Rahman, Dong-Hyun Kim

## Abstract

We present here a novel surface mass spectrometry strategy to perform untargeted metabolite profiling of formalin-fixed paraffin-embedded (FFPE) pediatric ependymoma archives. Sequential Orbitrap secondary ion mass spectrometry (3D OrbiSIMS) and liquid extraction surface analysis-tandem MS (LESA-MS/MS) permitted the detection of 887 metabolites (163 chemical classes) from pediatric ependymoma tumor tissue microarrays (diameter <1 mm; thickness: 4 μm). From these 163 classes, 60 classes were detected with both techniques, whilst LESA-MS/MS and 3D OrbiSIMS individually allowed the detection of another 83 and 20 unique metabolite classes, respectively. Through data fusion and multivariate analysis, we were able to identify key metabolites and corresponding pathways predictive of tumor relapse which were retrospectively confirmed using gene expression analysis with publicly available data. Altogether, this sequential mass spectrometry strategy has shown to be a versatile tool to perform high throughput metabolite profiling on sample-limited tissue archives.

For Table of Contents Only

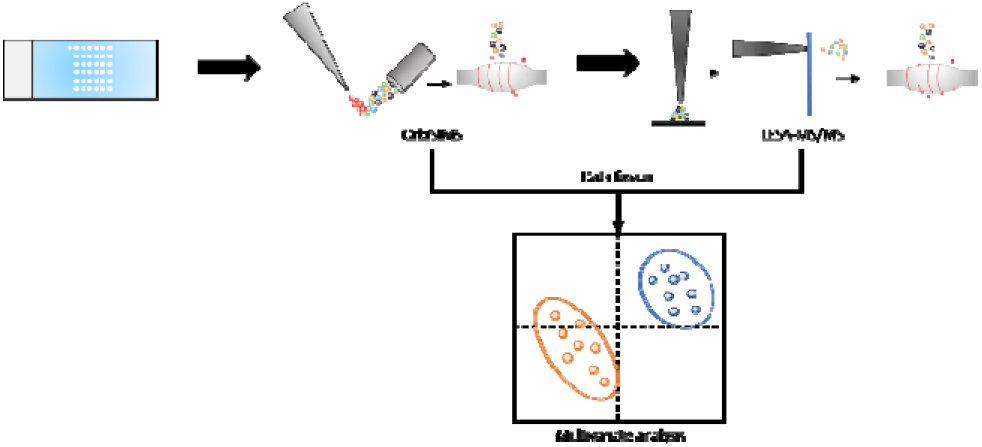

---

Central nervous system (CNS) pediatric tumors are the most prevalent type of solid cancer diagnosed in children and the leading cause of mortality among all cancers in children.^1^ From the clinical and biological perspective, intracranial pediatric ependymomas remain enigmatic and challenging tumors to treat. Overall, the prognosis is poor with over 50% of tumors relapsing and less than 50% of children surviving this disease (5-year overall survival is ~25% for patients who have relapsed).^2–4^ Though widely considered a ‘surgical disease’, a significant proportion of patients experience relapse, even following complete surgical resection of the tumor. The identification of biological correlates of disease progression and patient-tailored therapeutic targets therefore remains a significant challenge in this disease. Understanding the biochemical nature of tumor development is of vital importance for the development of the next generation of treatments.^2^ In a disease state, the human metabolome is affected by several factors and therefore provides an excellent source of information to investigate disease-related alterations in metabolism.^5^ To do so, an untargeted metabolomics approach can be used to study molecular changes within and between tissue samples of different phenotypes.^6,7^ State-of-the-art metabolomics techniques allow the detection of hundreds to thousands of metabolites in a biological sample ^8^, where untargeted metabolomics of cancer tissue is undertaken using liquid chromatography-mass spectrometry (LC-MS) and gas chromatography-MS (GC-MS).^9,10^ Chromatography-based strategies allow the identification of a vast number of metabolites; however, these require 20-50 mg of tissue for metabolomics analysis.

Preserving tumor regions of interest is commonly achieved using the tissue microarray (TMA) format. The neuropathologist identifies and cuts out key regions in the whole tissue section which are stored as a separate formalin-fixed paraffin-embedded (FFPE) block. The TMA platform allows small amounts of tissue to be used for transcriptomic and histological analysis ^11–13^, though for diagnosis the entirety of the tumor needs to be reviewed. Since the TMA tissue sections are small (diameter <1 mm; thickness: 4 μm), sensitive analytical techniques are required for metabolomic studies to detect low-abundance metabolites. Despite the vast amount of available TMA libraries, metabolite profiling of tumor TMAs has been an unexplored territory due to incompatibility of LC-MS or GC-MS analysis. To perform MS analysis on TMAs, a sensitive technique is required to directly obtain a wide range of metabolites from small tissue sections.

A few studies have shown the potential of liquid extraction surface analysis-MS (LESA-MS) for untargeted metabolomics across a range of sample types. With LESAMS, liquid microjunction-based extraction can be performed on a flat sample surface to obtain the analytes of interest which are directly injected into the mass spectrometer.^14^ Hall *et al.* ^15^ found significantly changed profiles of lipids in non-alcoholic fatty liver disease (NAFLD) tissue using LESA-MS. This allowed discrimination between different stages of stearosis. Ellis *et al.* ^16^ used LESA-MS And for the analysis of single-cell arrays and could distinguish cell types based on lipid profiles showing the capability of performing single-cell metabolomics with LESA. We have also successfully used LESA-MS to identify metabolic changes in small volumes of urine samples from an intervention study.^17^ Basu *et al.* ^18^ used LESA-MS for direct metabolite profiling for several different breast cancer cell lines. The capability of LESA was shown to allow direct analysis of adherent cells with minimal sample preparation. Collectively, these studies have shown that from a limited amount of sample, metabolic changes could be measured accurately.

The recent development of Orbitrap secondary ion MS (3D OrbiSIMS) revealed new possibilities for metabolic profiling due to its capability of high mass accuracy and mass resolving power (>240,000 at m/z 200) at subcellular spatial resolution.^19^ In that study, it was shown that 3D OrbiSIMS can be used for 2D and 3D imaging of neurotransmitters, in situ identification of lipid species using tandem MS, and performing metabolomics profiling of single cells.^19^ One unmet scientific challenge for brain tumor research is the capability to perform metabolomics analysis on archived TMAs to understand tumor development and find potential targets for therapies.^8,20^

3D OrbiSIMS and LESA tandem MS (LESA-MS/MS) require only minimal sample preparation, analysis can be performed directly on the tissue sample, and both instruments can acquire data in an automated manner using the TMA as sample platform. These MS techniques can therefore circumvent the need for tissue homogenization, allowing the tissue to remain architecturally intact and available for subsequent study. The data achievable using this approach will address current challenges in cancer metabolomics, as detection of low abundance (highly polar) oncometabolites to study important metabolic pathways may enable the development of novel prognostic and treatment strategies.^9,21^ To date, no disease studies have thus far reported the use of combined 3D OrbiSIMS and LESA-MS/MS for untargeted metabolite profiling on TMAs. Combining MS techniques, in which ions are generated via different mechanisms, will allow acquisition of complementary metabolomic datasets from the same set of samples. Combination of the individual datasets on existing TMA archives could therefore provide a vast amount of clinically valuable information. Here, we perform 3D OrbiSIMS and LESA-MS/MS analysis of FFPE pediatric ependymoma TMAs as an exemplar demonstration of the ability to perform untargeted surface metabolomics obtain clinically relevant data.

## EXPERIMENTAL SECTION

Tissue microarray preparation. Hematoxylin and eosin-stained sections from FFPE pediatric ependymoma (collection period 1992-2010) were examined by a neuropathologist at Nottingham University Hospital, and three representative areas were marked on the slides. Using a Raymond Lamb tissue micro-arrayer, 1 mm cores were punched from the marked areas of the donor blocks and placed into recipient paraffin blocks to generate a tissue microarray (TMA) on glass slides. 4-μm sections were cut from each block for use in further experiments. The analyzed TMA blocks consisted of patients who experienced tumor relapse (N = 5; n =3) and patients without relapse (N = 2; n = 3). All tissue sections were approximately 1 mm in diameter.

### Sample preparation for MS analysis

Deparaffinization of FFPE pediatric ependymoma TMAs (Fig. 1) was achieved using an adapted protocol from Ly *et al.*^22^ FFPE TMAs were first washed twice for 1 minute in a xylene bath (mixture of isomers; ≥98.5%, AnalaR NORMAPUR®, VWR, Leicestershire, UK). Residual xylene was removed, and the array was allowed to dry in a fume hood for at least one hour, before storage at room temperature until analysis. Time between storage and 3D OrbiSIMS analysis was less than 24 hours.

**Figure 1:**
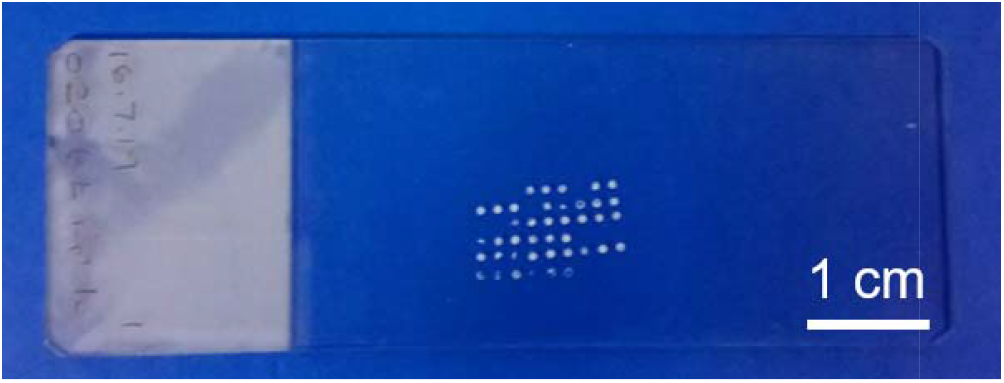
Example of a ependymoma tissue microarray before and after paraffin removal with xylene

### 3D OrbiSIMS

The TMA was placed in a hybrid TOF.SIMS 5 (IONTOF GmbH, Münster, DE) instrument coupled to a Q Exactive HF (Thermo Scientific, San Jose, CA) mass spectrometer without any dessication. Ions were sputtered from the surface using a 20 keV Ar_3000_^+^ gas cluster ion beam (GCIB). The field of view (FoV) was set to 500 μm × 500 μm around the centre of each tissue section to avoid charging effects of the glass surface. The ion dose was 6.3 × 10^14^ ions/cm^2^. Spectra were acquired at a lateral resolution of 20 μm in random raster mode. The Orbitrap™ was operated in Full-MS mode. The resolution was set to 240,000 at *m/z* 200 and the AGC target was set to 1 × 10^6^ with a maximum ion injection of 511 ms. Data were acquired in the scan range *m/z* 75-1125 for both positive and negative polarity.

### LESA-MS/MS

The TMA was placed on a universal plate holder (Advion Biosciences, Ithaca, NY) and scanned with an Epson V330 scanner. The tissue sample location was selected in LESA Points (Advion Biosciences, Ithaca, NY). Liquid extraction surface analysis-tandem mass spectrometry (LESA-MS/MS) was carried out using a TriVersa Nanomate (Advion Biosciences, Ithaca, NY) coupled to a Q Exactive plus Orbitrap mass spectrometer (Thermo Scientific, San Jose, CA). Extraction of metabolites from tissue samples was conducted with a mixture of 80% v/v methanol (CHROMASOLV®; Sigma-Aldrich, Gillingham, UK), and 20% v/v water (CHROMASOLV®; Sigma-Aldrich, Gillingham, UK) to which MS-grade formic acid (Optima™ LC-MS grade; Fisher Scientific, Loughborough, UK) was added (final concentration 1% v/v). Brain tissue was sampled using the contact LESA approach^23^ in which the solvent tip is brought into contact with the sample to minimize the solvent spread. During contact, 1.5 μL solvent (total volume: 3 μL) was dispensed on the tissue and after 15 seconds, 2.0 μL was aspirated back into the tip. The extract was introduced into the mass spectrometer via chip-based nanoelectrospray ionization (ESI Chip™, Advion Biosciences, Ithaca, NY) at 1.4 kV and 0.3 psi gas pressure.^17^ The mass spectrometer was operated in Full-MS/dd-MS^2^ mode. MS^1^ spectra were acquired in the scan range of *m/z* 70-1050. The resolution was set to 140,000 at *m/z* 200 and the AGC target was set to 3 × 10^6^ with a maximum ion injection time of 200 ms. Data-dependent MS/MS spectra were acquired at a resolution of 17,500 at *m/z* 200. The AGC target for MS^2^ scans was set to 1 × 10^5^ with a maximum ion injection time of 50 ms. The top 20 most intense ions were isolated within a 1 *m/z* window for fragmentation. Dynamic exclusion was applied for 120 seconds per polarity. Fragmentation was carried out by higher-energy collisional dissociation (HCD) using a stepped collision energy of respectively 10, 25 and 40 eV. All tissue sections were analyzed once. MS data were acquired for 2 minutes per polarity. Time between 3D OrbiSIMS and LESA-MS/MS analysis was less than 24 hours.

### Feature extraction

Mass spectrometry data were processed using an in-house MATLAB (R2017a, The MathWorks, Inc., Natick, MA) script. For LESA-MS data, files were converted to. mzXML using ProteoWizard (v3.0.1908).^24^ Peaks were picked from averaged spectra using the mspeaks function (threshold: 1% of base peak intensity) and aligned within 5 ppm m/z window.^15^ Features with >20% missing values across all samples were removed.^25^ Remaining missing values were imputed using *k*-nearest neighbor (*k*nn) imputation. The value of *k* was set to 10.^26^ For 3D OrbiSIMS, total ion spectra were exported as. TXT files and further processed in MATLAB as outlined above. The threshold intensity for peaks was set to 1% of the base peak intensity. The output intensity matrices were stored in. XLSX format for further use.

### Metabolite identification and pathway analysis

Peak lists were searched against the Human Metabolome Database^27^ with 3 ppm mass tolerance using [M+H]^+^, [M+Na]^+^, [M+K]^+^, and [M+H-H_2_O]^+^ as ions for positive mode and [M-H]^-^, and [M-H-H_2_O]^-^ for negative mode. Monoisotopic masses were exported from the Human Metabolome Database and then submitted to MetExplore^28^ for pathway analysis. Masses were searched against the Homo sapiens (Strain: global) (Source: Publication, Version: 2.02) database (3 ppm mass tolerance).

### Data fusion and statistical analysis

Analyzed ependymoma samples were divided in two groups: *no relapse* and *eventual relapse* for patients who did not experience relapse and patients for whom ependymoma relapsed after surgical removal. For data fusion, a low-level strategy^29^ was used in which the individual ion intensity matrices derived from the feature extraction workflow were normalized to the total ion count (TIC) and log-transformed and subsequently concatenated. Next, data were subjected to partial least squares-discriminant analysis (PLS-DA). An initial model was built for feature selection based on a variable’s importance in projection (VIP) score ≥ 1.5. With the selected variables, a new PLS-DA classification model was created and validated through leave-one-out cross validation and a permutation test.^30^ Leave-one-out cross validation was performed by holding out all replicates for one subject during each iteration.

Univariate statistical analysis was Student’s *t*-test. False discovery rates were estimated using permutations. A p-value < 0.05 was considered significant.

## Results & Discussion

### The sequential mass spectrometry workflow for untargeted metabolomics

We analyzed de-paraffinized TMAs (Fig. 2A-B), first with 3D OrbiSIMS (Fig. 2C), followed by LESA-MS/MS (Fig. 2D). The idea behind this analysis strategy was to maximize the metabolite coverage whilst consuming only a minimal amount of tissue using complementary MS techniques. After the data were acquired, they were processed in MATLAB for peak detection and alignment (Fig. 2E). To get the most out of the 3D OrbiSIMS and LESA-MS/MS datasets, a low-level data fusion strategy^25,29^ was added to the workflow, which means that the ion intensity data from 3D OrbiSIMS and LESA-MS/MS were combined into one single data matrix instead of creating individual classification models for each dataset (Fig. 2F). The fused data was then subjected to PLS-DA with subsequent permutation testing^30^ to select discriminative ions between ependymoma subgroups (Fig. 2G). Molecular formulae were subsequently assigned using the Human Metabolome Database^27^ (Fig. 2H) and submitted for pathways analysis in MetExplore^28,31^ to identify significantly affected metabolic pathways and corresponding genes (Fig. 2I).

**Figure 2:**
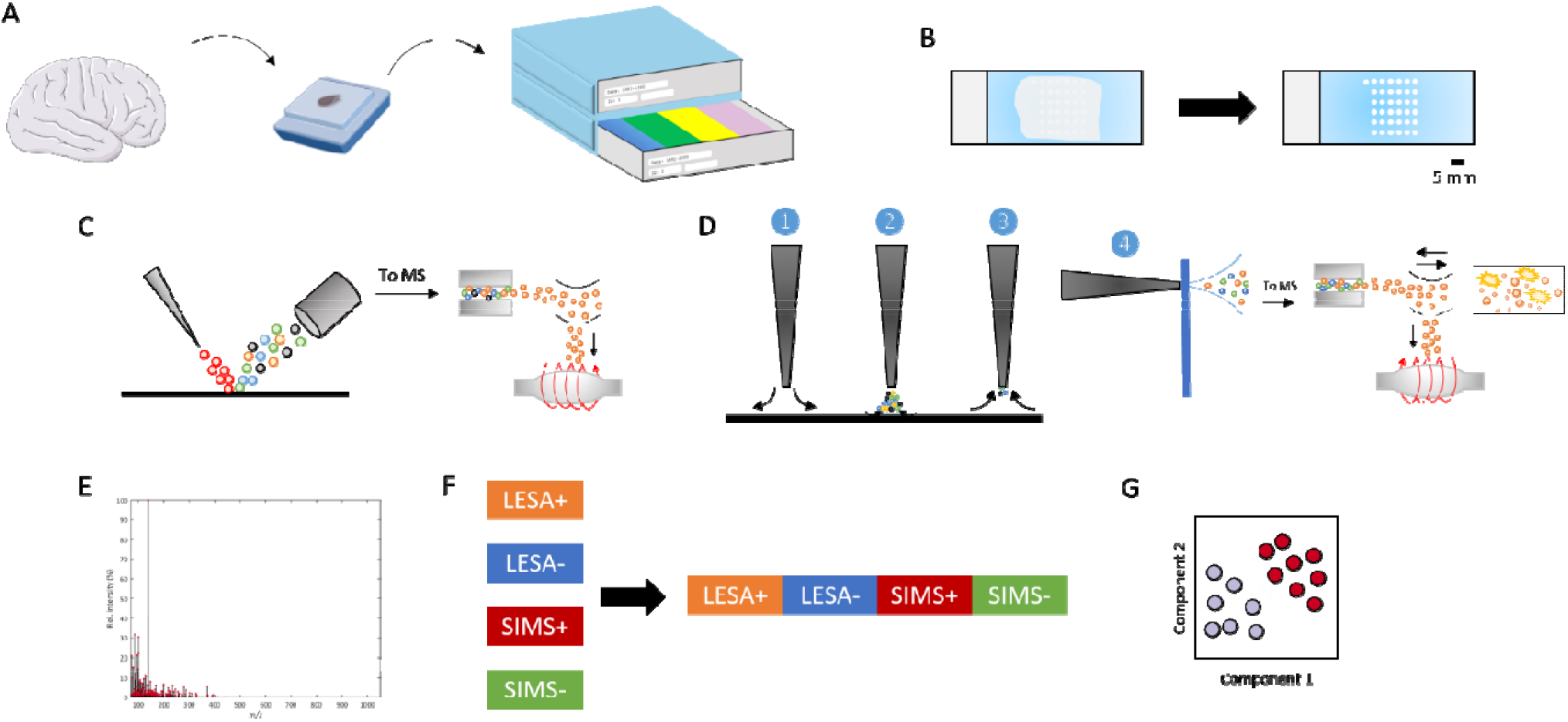
Sequential mass spectrometry analysis of pediatric ependymoma tissue microarrays. (A) The tumor tissue was removed, the tumor area was marked and then paraffin-embedded for long term storage. (B) For MS analysis, a TMA block from the archive was sectioned and mounted onto a glass substrate followed by a xylene wash to remove the paraffin. (C) Paraffin-free samples were then analyzed by OrbiSIMS followed by (D) LESA-MS/MS. (E) Ions were selected from the mass spectra and aligned. (F) All matrices with ion intensities were then combined (low-level data fusion). (G) Subsequently, data were subjected to partial-least squares-discriminant analysis (PLS-DA) to identify discriminative features in tumor recurrence. Molecular formulae were assigned to the significant ions using the Human Metabolome Database. Ions with a putative ID were then submitted to MetExplore for metabolic pathway analysis to identify affected pathways and corresponding genes.

### Complementary metabolite profiling with 3D OrbiSIMS and LESA-MS/MS

In both MS techniques, molecules are ionized via different mechanisms. This would potentially allow coverages of a wider range of metabolites since molecules might be more efficiently ionized with either technique. Representative mass spectra for each instrument are shown in Figure S1. During SIMS analysis, the sample was slightly etched by the primary ion beam (20 keV Ar_3000_^+^), although the amount of sample consumed by SIMS was limited when argon clusters were used.^32^ Prior 3D OrbiSIMS analysis did neither deplete the ion intensities (Student’s *t*-test: p = 0.8345; Figure S2) nor reduce the number of features (Student’s *t*-test: p = 0.4743; Figure S3) in subsequent LESA-MS/MS analysis. In total, 634 and 51 ions were assigned a molecular formula for LESA-MS data acquired in positive and negative ionization modes, respectively. For 3D OrbiSIMS data, 86 and 116 ions were annotated with putative IDs in positive and negative ionization modes, respectively (Fig. 3A). The number of identified metabolites is a vast improvement compared to previous research done on metabolite profiling of ependymoma tissue using nuclear magnetic resonance spectroscopy (NMR).^1,33^ Similar numbers of metabolites were identified using LC-MS(/MS) in different types of cancer FFPE tissue.^34–36^

**Figure 3:**
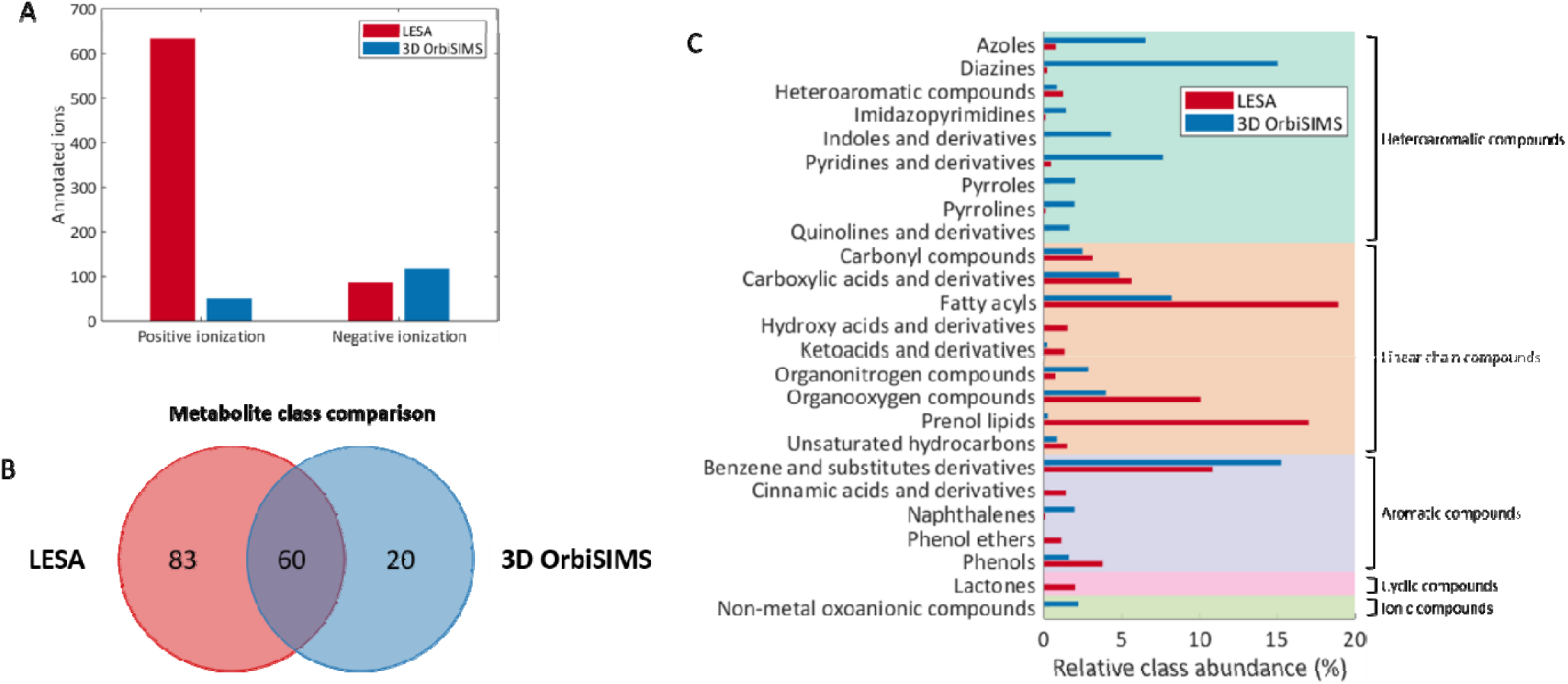
Identifying putative metabolite features in OrbiSIMS and LESA-MS spectra. (A) In total, more features were identified in the LESA-MS spectra, though the number of ions in negative mode identified as metabolite was higher for SIMS. (B) For all identified features, the class as described in the HMDB was obtained to identify which classes can be detected with either technique. The patched areas represent structurally similar classes. (C) From the Venn diagram can be derived with both surface mass spectrometry techniques unique metabolite classes can be detected and could therefore provide complementary information.

To assess the degree of complement in annotated metabolites, HMDB classes were derived from the putative identities. Twenty unique metabolite classes were found using 3D OrbiSIMS, followed by another 83 unique metabolite classes with LESA-MS/MS (Fig. 3B). Further investigation of metabolite class breakdown reveals that with SIMS predominantly non-polar metabolites can be detected whilst LESA-MS/MS permits the analysis of polar metabolites (Fig. 3C). This reveals the benefit of analyzing the same sample set with complementary MS techniques for increased metabolite coverage.

### Fused metabolite profiles reveal signatures predictive of brain tumor relapse

The analyzed sample cohort (N = 7; n = 3) consisted of primary pediatric ependymomas (Table 1). For 5 patients it was known that the tumor eventually recurred. To assess whether any alteration in the metabolite profile could be observed, patients were divided into no relapse (N = 2; n = 3) and eventual relapse groups (N = 5; n = 3). Through data fusion and multivariate analysis (partial least squares-discriminant analysis (PLS-DA)), we were able to cluster patients based on whether the tumor eventually recurred (Fig. 4A). To identify the discriminative ions between *no relapse* and *eventual relapse* ependymoma cohorts, the VIP score for each ion was calculated. A VIP score ≥ 1.5 was considered discriminative. After PLS-DA, the model was validated using leave-one-out cross validation which resulted in a Q^2^ (goodness-of-prediction) of 0.4606. The PLS-DA model showed an acceptable Q^2^ (>0.4) for a biological model.^37^ Through a permutation test^30^, it was shown that no random PLS-DA model predicted tumor recurrence better than the original PLS-DA model (p < 0.05).

**Table 1:** Demographic information for patients included in the analysis per class

| Class | Total patients | Age (months) | Gender | WHO grade | Ependymoma class | Tumor site | Ki-67 score | Nucleolin score |
| --- | --- | --- | --- | --- | --- | --- | --- | --- |
| Eventual relapse | 5 | 35 ± 25 | M: 1/ F: 4 | II: 3<br>III: 2 | Anaplastic: 2<br>NOS: 3 | PF: 5 | < 1: 2<br>1: 2<br>N/A: 2 | 95: 2<br>65: 1<br>N/A: 2 |
| No relapse | 2 | 64 ± 31 | M: 1/ F: 1 | II: 0<br>III: 2 | Anaplastic: 2<br>NOS: 0 | PF: 1<br>ST: 1 | N/A: 2 | N/A: 2 |

NOS: not otherwise specified; PF: posterior fossa; ST: supratentorial

**Figure 4:**
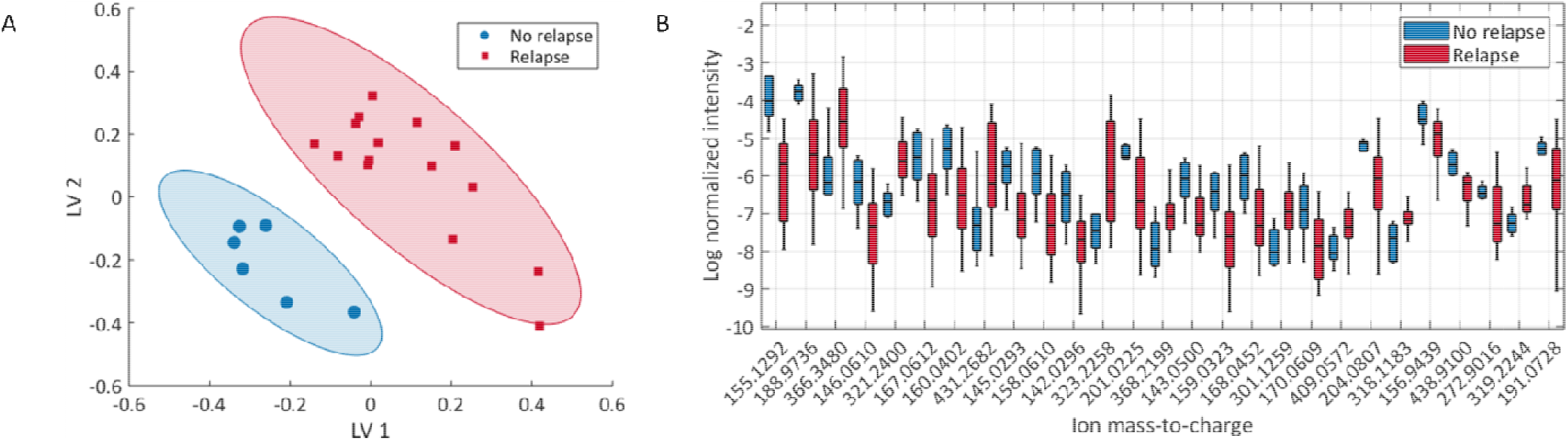
Identification of significant ions from 3D OrbiSIMS and LESA-MS/MS data. (A) PLS-DA scores plot after data fusion reveals clustering of patients based on tumor recurrence. (B) Box plot for significant ions (p < 0.05) identified using Student’s t-test and FDR estimation through a permutation test.

The ions which met this criterion were subjected to Student’s *t*-test to determine which ions were significantly altered between the two groups. In total, we identified 27 significant mass ions in the fused data set (p < 0.05; Fig. 3B). Classification of ependymoma subgroups substantially using significant ions only (Q^2^: 0.6375).

From the significant 27 mass ions, 18 mass ions were assigned putative molecular formulae using the Human Metabolome Database^27^ (Table 2). From those 18 ions, 6 were detected with LESA-MS and the other 12 ions with 3D OrbiSIMS. For all significant ions, the fold change in ion intensity was calculated from the fused data matrix between the *no relapse* and *eventual relapse* groups. Most of the significant ions were found to be more prominent in the no relapse group.

**Table 2:** Annotations for significant ions (p < 0.05). Fold changes were calculated by dividing the average ion intensity of the *no relapse* group by the average ion intensity in the *eventual relapse* group

| $m/z$ | Formula | $\Delta ppm$ | Adduct | Instrument | Fold change |
| --- | --- | --- | --- | --- | --- |
| 321.2400 | $C_{18}H_{34}O_3$ | 0 | $[M+Na]^+$ | LESA | 0.23 |
| 145.0293 | $C_9H_6O_2$ | 2.1 | $[M+H-H_2O]^+$ | LESA | 1.98 |
| 201.0225 | $C_8H_{10}O_5S$ | 1.5 | $[M+Na]^+$ | LESA | 1.64 |
| 143.0500 | $C_5H_{12}OS$ | 0.7 | $[M+Na]^+$ | LESA | 1.57 |
| 170.0609 | $C_{11}H_9NO_2$ | 2.4 | $[M+H-H_2O]^+$ | LESA | 1.89 |
| 319.2244 | $C_{18}H_{32}O_3$ | 0 | $[M+Na]^+$ | LESA | 0.45 |
| 146.0610 | $C_9H_{11}NO_2$ | 2.7 | $[M-H-H_2O]^-$ | SIMS | 2.53 |
| 167.0612 | $C_{11}H_8N_2$ | 1.8 | $[M-H]^-$ | SIMS | 2.42 |
| 160.0402 | $C_9H_9NO_3$ | 1.9 | $[M-H-H_2O]^-$ | SIMS | 2.39 |
| 145.0293 | $C_9H_6O_2$ | 1.4 | $[M-H]^-$ | SIMS | 1.98 |
| 158.0610 | $C_{10}H_{11}NO_2$ | 2.5 | $[M-H-H_2O]^-$ | SIMS | 2.25 |
| 142.0296 | $C_9H_7NO_2$ | 2.1 | $[M-H-H_2O]^-$ | SIMS | 1.57 |
| 201.0225 | $C_8H_{10}O_4S$ | 1.0 | $[M-H]^-$ | SIMS | 1.64 |
| 143.0500 | $C_{10}H_8O$ | 1.4 | $[M-H]^-$ | SIMS | 2.50 |
| 168.0452 | $C_{11}H_9NO_2$ | 1.8 | $[M-H-H_2O]^-$ | SIMS | 1.94 |
| 170.0609 | $C_{11}H_{11}NO_2$ | 1.8 | $[M-H-H_2O]^-$ | SIMS | 1.89 |
| 409.0570 | $C_{21}H_{14}O_9$ | 1.2 | $[M-H]^-$ | SIMS | 0.44 |
| 318.1183 | $C_{13}H_{32}NO_9$ | 1.9 | $[M-H-H_2O]^-$ | SIMS | 0.49 |

The same processing workflow was used to determine the benefit of data fusion. The same ions were identified as being discriminative between no relapse and eventual relapse groups. Data fusion resulted in a slightly higher Q^2^ value mode compared to the individual data sets per MS method (Table S1). The data fusion workflow allows processing four individual datasets at once and resulting in a single classification model is generated, making this workflow more efficient. Also, this workflow is not only applicable for LESA-MS and 3D OrbiSIMS data but could be used for any type of mass spectrometry data.

Previous studies (e.g. Mascini *et al.*^38^) have shown that tumor classification could be done successfully using matrix-assisted laser desorption/ionization-mass spectrometry imaging (MALDI-MSI). The results here show that combined or single LESA and 3D OrbiSIMS can be used as alternative for tumor classification. The advantage of LESA and 3D OrbiSIMS is that the required sample preparation only consists of paraffin removal (i.e. no matrix has to be applied on to the TMA).

We further investigate the significant ions obtained using 3D OrbiSIMS. Due to the hard ionization process in SIMS, molecules tend to fragment. Therefore, putative assigned IDs could potentially be an isobaric fragment. Ion intensities were pairwise compared using Pearson’s correlation coefficient (Figure S4). Ions with a correlation > 0.95 were considered to belong to the same parent/fragment. We found that none of the significant ions had a strong correlation with another ion in the 3D OrbiSIMS data (Table S2) and therefore it suggests that there is no contribution to these ions from other fragment/parent ions.

The putative metabolite IDs were submitted to MetExplore^28,31^ for metabolic pathway analysis. This allowed putative identification of affected pathways between ependymoma subgroups. We could identify five metabolic pathways with 14 associated genes to be potentially affected between the *no relapse* and *eventual relapse* groups (Table 3). To the best of our knowledge, this is the first report of investigating alterations in MS-based metabolite profiles between ependymoma subgroups. Previous work on metabolite profiling of ependymoma via NMR showed that L-phenylalanine is highly abundant and an important discriminative metabolite for ependymoma among other pediatric brain tumors.^1,33^ Here, we found a more prominent abundance of L-phenylalanine (*m/z* 146.0610; fold change: 2.53) in the no relapse group. We also putatively identified metabolites in the tryptophan metabolism pathway (5-hydroxytryptophol, 4,6-dihydroxyquinoline, β-carboline, methyl indole-3-acetate and quinolone-4,8,-diol) to be significantly increased between the no relapse and eventual relapse ependymoma cohorts. Previous work has identified phenylalanine and tryptophan metabolism to be important pathways in brain tumor metabolism.^9^ Cytochrome metabolism and tyrosine metabolism were also found to be increased significantly in the no relapse group. Also, these pathways have not been reported concerning ependymoma relapse. Cytochrome enzymes are important in the metabolism of endogenous and exogenous compounds and could be important factors in drug therapy resistance.^39,40^ Tyrosine metabolism pathways have been proposed as potential targets for treatment of glioblastoma.^41^ Conversely, linoleate metabolism was more predominant in the eventual relapse group for the conversion of epoxy fatty acids (EpOMEs) to fatty acids and vice versa. Whilst linoleate metabolism has not been reported for ependymoma relapse, fatty acid oxidation pathways have been reported to be affected in carcinogenesis.^42,43^ These observations indicate that the sequential MS strategy permits the identification of important metabolic changes associated with brain tumor progression from small amounts of tissue in a high-throughput manner.

**Table 3:** Putative metabolite IDs for significantly affected pathways (p < 0.05) and related genes between ependymomas which relapsed and those which did not relapse

| Pathway | Putative metabolite ID | Related genes |
| --- | --- | --- |
| Tryptophan metabolism | 4,6-dihydroxyquinoline<sup>S</sup> (↑) | ADHFE1<br>MAOA<br>MAOB |
| | 5-hydroxytryptophol<sup>S</sup> (↑) | |
| | β-carboline<sup>S</sup> (↑) | |
| | Methyl indole-3-acetate<sup>S</sup> (↑) | |
| | 4,8-dihydroxyquinolone<sup>S</sup> (↑) | |
| Linoleate metabolism | 12,13-EpOME<sup>L</sup> (↓) | EPHX1<br>EPHX2 |
| | 9,10-EpOME<sup>L</sup> (↓) | |
| Cytochrome metabolism | Coumarin<sup>S</sup> (↑) | CYP2A13<br>CYP2A6<br>CYP2F1 |
| | Napthalene epoxide<sup>S</sup> (↑) | |
| Phenylalanine metabolism | L-phenylalanine<sup>S</sup> (↑) | DDC |
|  |  | GOT1 |
|  |  | GOT2 |
|  |  | PAH<br>TAT |
| Tyrosine metabolism | Adrenochrome<sup>S</sup> (↑) | GSTK1 |

Arrows indicate higher ( ↑ ) or lower ( ↓ ) abundance of the metabolite in the *eventual relapse* group compared to the *no relapse* group. <sup>L</sup>: identified using LESA-MS. <sup>S</sup>: identified using 3D OrbiSIMS.

### Validation of MS-based metabolomics with publicly available gene expression data

Due to the limited availability of ependymoma samples, only a small cohort could be analyzed. We recognize the need for an increase in the number of patients to validate these results, though the data shows the clinical potential for the sequential MS strategy to be further used for metabolomics screening of ependymoma TMAs. To validate our current findings, we used publicly available gene expression datasets for primary (n = 72) and recurrent (n = 47) pediatric ependymoma.^44,45^ Ten out of the fourteen genes listed in Table 3 were present in the gene expression datasets. From those 10 genes, four genes showed a significant differential expression. ADHFE1 (p = 0.00234) was upregulated in primary ependymoma, whilst GSTK1 (p = 0.00450), GOT2 (p = 0.0393) and EPHX2 (p = 0.0433) showed a significantly higher expression in the recurrent ependymoma cohort. The gene expression results are partially in concordance with the metabolomics data. The expression of GSTK1 and GOT2 is in line with higher abundance of adrenochrome, and L-phenylalanine, respectively. On the other hand, the expression of ADHFE1 and EPHX2 was opposite to the observed difference in metabolite abundance between the no relapse and eventual relapse group. An explanation could be that for the metabolomics study only primary ependymomas were used and these might not completely reflect the same gene expression profile as recurrent ependymoma and would require further investigation using a larger sample cohort. Also, the group of ten genes as a single gene set showed significant change between primary and recurrent ependymoma (p = 0.0 257, global test) and showed clear separation of primary tumors from recurrent ependymal tumors (Fig. 5), indicating a good predictive power of the four significant genes for predicting ependymoma relapse. Although our metabolomics dataset is small, the gene expression analysis supports the significance of the genes we identified through MS-based metabolomics and pathway analysis. Moreover, excellent clustering was observed for primary and recurrent ependymoma based on the subset of the significant genes. This confirmation of our metabolomics data shows the potential of the sequential MS strategy to be further used for large scale clinical studies on archived TMAs.

**Figure 5:**
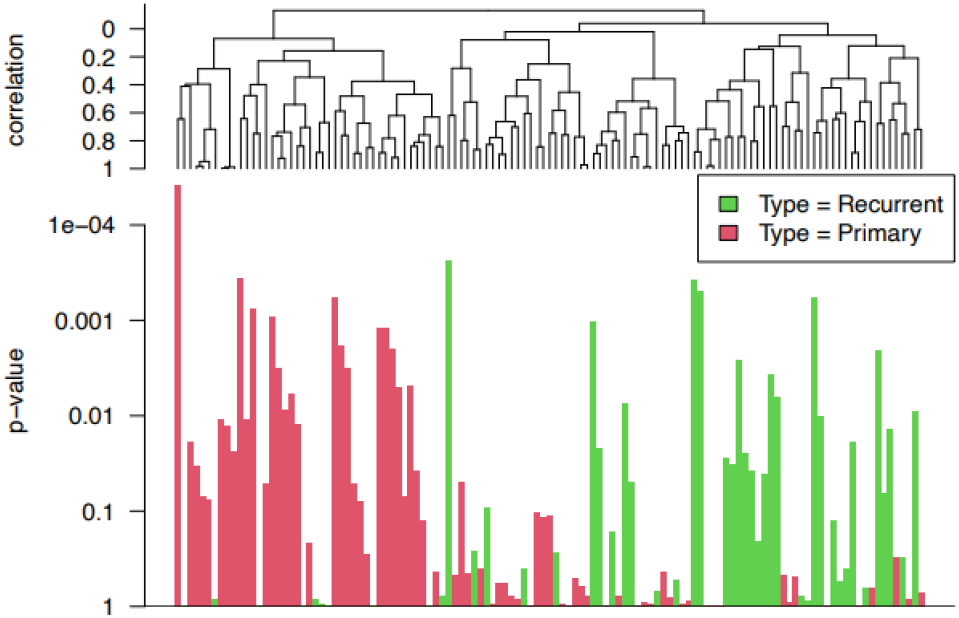
Gene expression analysis of primary (n = 72) and recurrent ependymoma (n = 47). The dendrogram reveals two distinct groups for which most of the recurrent samples belong to the first group and most of the primary samples to the second group. The samples associated with strong evidence for the association between the response (primary vs recurrent) and the gene expression profile of the gene set (10 genes) have small p-values (tall bars in the bottom plot)

## Conclusion

We have presented a novel mass spectrometry strategy for metabolite profiling of tumor microarrays. Complementary metabolite profiles were obtained permitting putative identification of additional affected metabolic pathways and their corresponding genes, resulting in a putative predictive signature of no relapse/relapse. This opens new opportunities to perform large scale metabolomics studies on archived tissue libraries. Furthermore, the minimally required sample preparation and short analysis time (10 minutes with 3D OrbiSIMS; 4 minutes with LESA-MS/MS) permits high sample throughput, making this strategy a competitive alternative to standard metabolomics analysis such as GC-MS, LC-MS and NMR.

## ASSOCIATED CONTENT

**Supporting Information**

**S1:** Statistical analysis of the effect of 3D OrbiSIMS analysis on subsequent LESA-MS analysis

**S2:** Representative mass spectra for ependymoma tissue sections acquired with 3D OrbiSIMS and LESA-MS/MS

**S3:** Multivariate analysis of individual LESA-MS and 3D OrbiSIMS data sets

**S4:** Ion correlation for 3D OrbiSIMS spectra in negative ionization mode Mass spectrometry data (3D OrbiSIMS and LESA-MS/MS) is available from the Nottingham Research Data Repository (https://rdmc.nottingham.ac.uk/)

## AUTHOR INFORMATION

### Author Contributions

The manuscript was written through the contributions of all authors. All authors have approved the final version of the manuscript.

## ACKNOWLEDGMENT

This work was supported by the Engineering and Physical Sciences Research Council [grant number: EP/N006615/1]; grant name: EPSRC Programme Grant for Next Generation Biomaterials Discovery

