## Supplementary Information for "Sequential 3D OrbiSIMS and LESA-MS/MS-based metabolomics for prediction of brain tumor relapse from sample-limited primary tissue archives"

**Supporting information**

**S1:** Representative mass spectra for ependymoma tissue sections acquired with 3D OrbiSIMS and LESA-MS/MS

**S2:** Statistical analysis of the effect of 3D OrbiSIMS analysis on subsequent LESA-MS analysis

**S3:** Multivariate analysis of individual LESA-MS and 3D OrbiSIMS data sets

**S4:** Ion correlation for 3D OrbiSIMS spectra in negative ionization mode

**S1:** Representative mass spectra for ependymoma tissue sections acquired with 3D OrbiSIMS and LESA-MS/MS

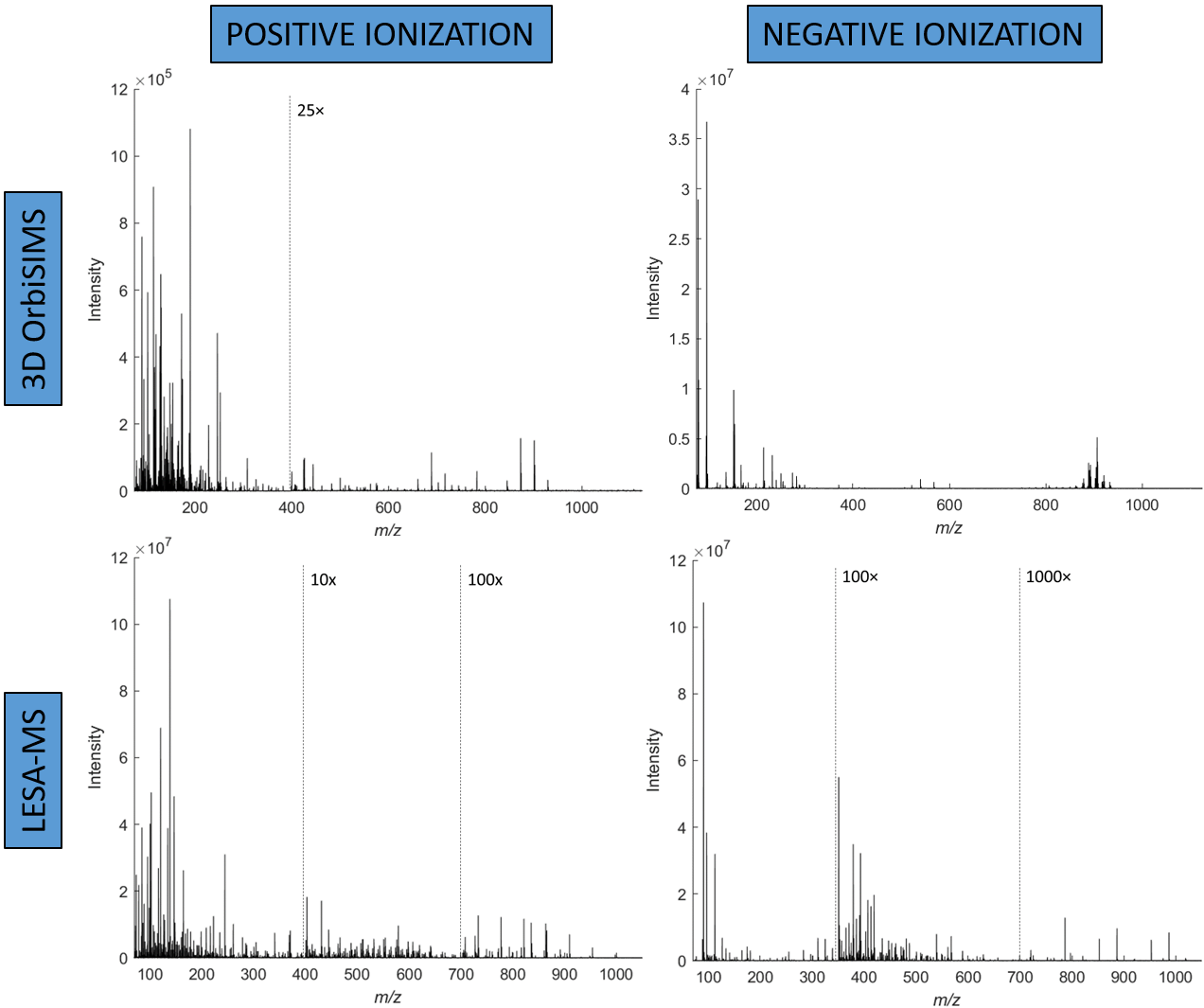

Figure S1: Representative mass spectra for ependymoma tissue sections acquired with 3D OrbiSIMS and LESA-MS/MS

**S2:** Statistical analysis of the effect of 3D OrbiSIMS analysis on subsequent LESA-MS/MS analysis

Tissue sections from one patient were chosen to test the effect of 3D OrbiSIMS analysis on subsequent LESA-MS/MS analysis. Analysis was performed on three repeats with prior 3D OrbiSIMS followed by LESA-MS/MS and another three repeats were directly analyzed with LESA-MS/MS. It was found that 3D OrbiSIMS analysis did neither lead to a depletion of the ion intensities for subsequent LESA-MS/MS analysis nor did it reduce the number of detected features.

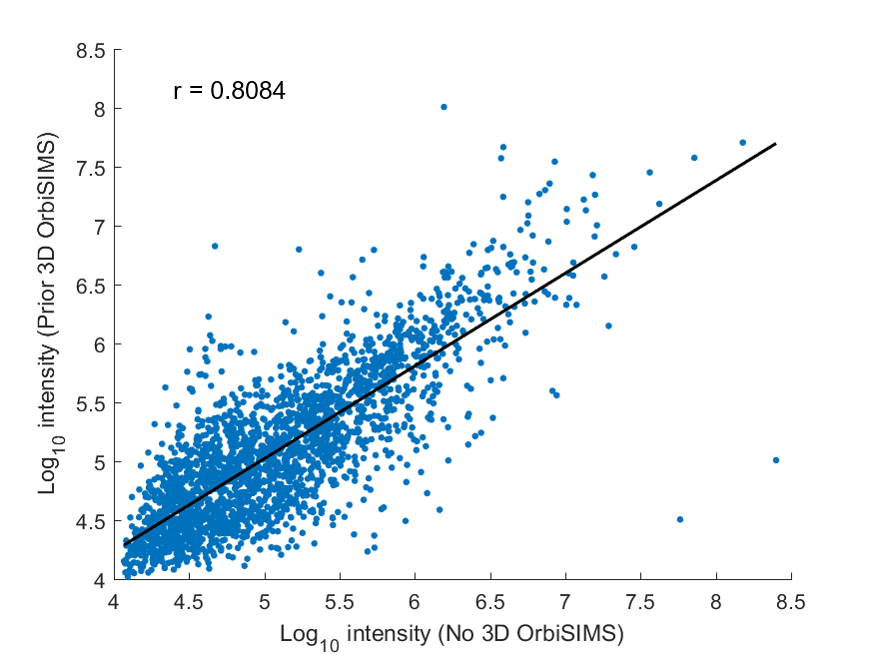

Figure S2: Distribution of metabolite ion signal intensities with and without prior 3D OrbiSIMS analysis.

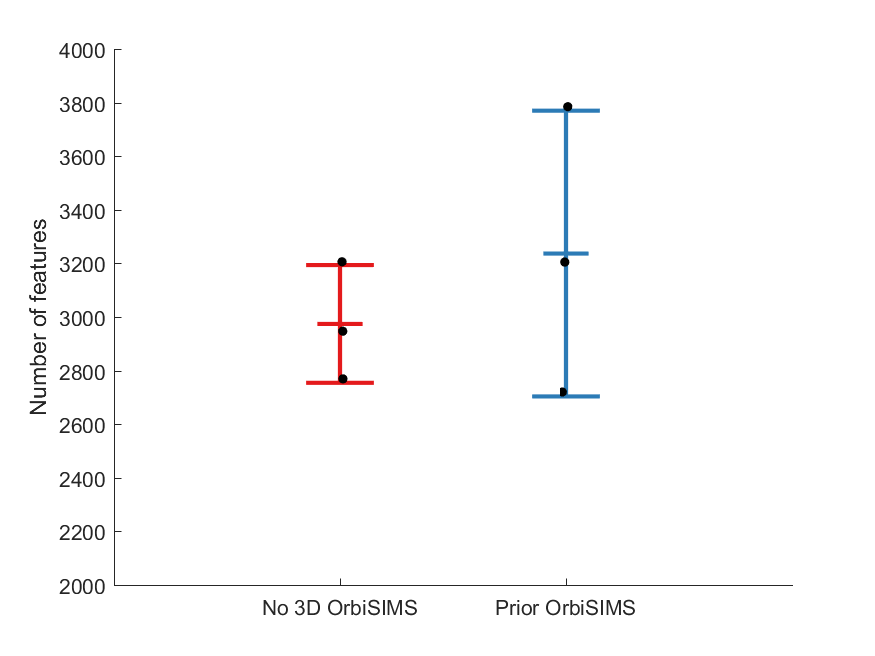

Figure S3: Comparison of number of LESA-MS features with and without prior 3D OrbiSIMS analysis. No difference in number of features was observed (p = 0.4743).

**S3:** Multivariate analysis of individual LESA-MS and 3D OrbiSIMS data sets

To compare multivariate analysis results from data fusion with separate multivariate analysis per peak matrix, LESA-MS and 3D OrbiSIMS data were individually subjected to PLS-DA. As can be seen, both LESA-MS data acquired in positive mode and 3D OrbiSIMS data acquired in negative mode are both able to classify no relapse and eventual relapse samples to the same degree as the fused data.

Table S1: PLS-DA classification of no relapse and eventual relapse ependymoma samples using individual datasets

| **Dataset** | **Q^2^** |
| --- | --- |
| LESA (+) | 0.4415 |
| LESA (-) | -0.2597 |
| SIMS (+) | -0.1741 |
| SIMS (-) | 0.4081 |

**S4:** Ion correlation for 3D OrbiSIMS spectra in negative ionization mode

Fragmentation of parent ions is commonly observed in SIMS spectra due to the hard ionization process. As can be seen in Supplementary Figure S2, many ions correlate well with each other. To investigate whether the putatively assigned ions are potentially derived from isobaric fragments, the correlation coefficients for the significant ions obtained from 3D OrbiSIMS analysis were investigated (Supplementary Table S2). None of the significant ions was found to have a strong correlation (r > 0.95) to other ions and therefore it suggests that there is no contribution to these ions from other fragment/parent ions.

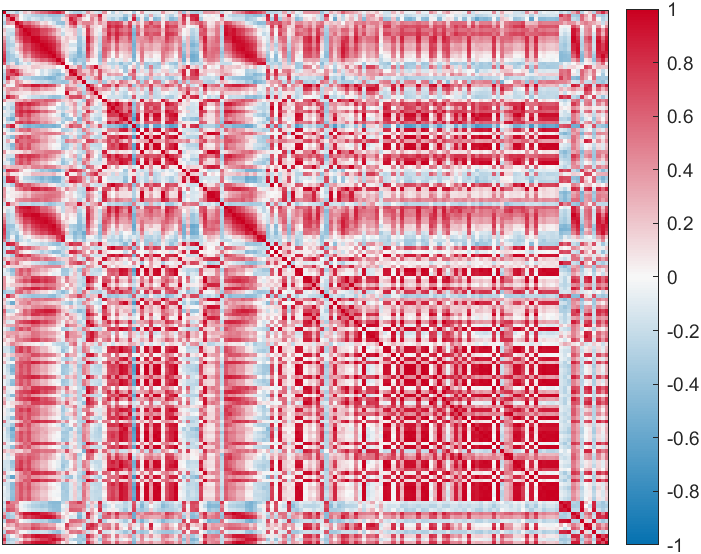

Figure S4: Correlation matrix for 3D OrbiSIMS ions obtained in negative ionization mode

Table S2: Overview of highly correlating ions with significant ions identified in the 3D OrbiSIMS spectra

| **m/z** | **Adduct** | **Correlating ions (r > 0.95)** |
| --- | --- | --- |
| 146.0610 | [M-H-H_2_O]^-^ | - |
| 167.0612 | [M-H]^-^ | - |
| 160.0402 | [M-H-H_2_O]^-^ | - |
| 145.0293 | [M-H]^-^ | - |
| 158.0610 | [M-H-H_2_O]^-^ | - |
| 142.0296 | [M-H-H_2_O]^-^ | - |
| 201.0225 | [M-H]^-^ | - |
| 143.0500 | [M-H]^-^ | - |
| 168.0452 | [M-H-H_2_O]^-^ | - |
| 170.0609 | [M-H-H_2_O]^-^ | - |
| 409.0570 | [M-H]^-^ | - |
| 318.1183 | [M-H-H_2_O]^-^ | - |
